# Hydroponics: A Novel Approach to Culturing *Aphanomyces euteiches*

**DOI:** 10.1101/2023.10.30.564802

**Authors:** Ethan Done

**Affiliations:** University of Saskatchewan

## Abstract

*Aphanomyces euteiches* is a soil-borne, economically impactful pathogen that afflicts pulse crops. Isolating it from environmental soil samples is made difficult by other microorganisms naturally present in field soil, particularly *Rhizopus*, that outcompete it on selective media. However, by isolating and culturing *A. euteiches* from pea plants grown in a hydroponics apparatus inoculated with field soil, this can be ameliorated to allow for enhanced observation and culturing of *A. euteiches* in media. This experiment tested and validated a method of hydroponic growth for *A. euteiches*, and confirmed that it yielded *A. euteiches* isolates using universal CPN60 sequencing.

## INTRODUCTION

*Aphanomyces euteiches* is a pathogen that affects legumes and pulse crops.^1^ Since it arrived in Saskatchewan in 2012, it has become a serious problem for producers.^1^ *A. euteiches* spores have also been found to have the ability to persist in infected field soil for up to ten years and have no known treatment.^1^ It is thus evident that isolating and culturing *A. euteiches* field isolates is of high importance to the agri-food sector. Various methods to isolate *A. euteiches* using infected root tissue placed on selective agar plates have long been developed. However, these can take up to two weeks to yield an isolate and often require further purification.^2,3^ This is due to many other microorganisms being co-selected with *A. euteiches* on selective plates: including *Rhizopus* and *Pythium*.^3^ Based on laboratory experience, *Rhizopus* is of particular concern, as it is visually identical to *A. euteiches* until after it must be isolated from the MVB plate and makes *A. euteiches* impossible to isolate once established due to its competitiveness.^3^ To eliminate these unwanted contaminants, soil known to contain *A. euteiches* was used to infect pea plants growing in a Kratky hydroponic system. The hypothesis is that this would dilute the *Rhizopus* while enriching *A. euteiches*, preventing it from outcompeting *A. euteiches* on the MVB plate. This method was refined using an iterative process until a protocol allowing for the successful isolation of *A. euteiches* from field soil was developed.

## MATERIALS AND METHODS

### Hydroponic Design

The hydroponic system was contained in a Neptune 1000ul filtered micropipette tip container (4.5 inches x 3.25 inches x 4 inches). The bottom of the container was covered in aluminum foil, and the sides were wrapped in VWR laboratory tape to prevent light from entering. A small closable window was made into the tape to allow for viewing of the roots. This created the base of the hydroponic system. The pipette tip holder tray was cut according to the design shown in Figure 1[A] to create the removable seed tray. Lastly, the top of the lid of the container was cut so that the pea plants could grow, and a strip of tape was added to reduce the amount of light that reached the seedlings, thus creating the upper chamber of the hydroponics system. Figure 1 [B] shows the completed setup after germination.

**Fig. 1.**
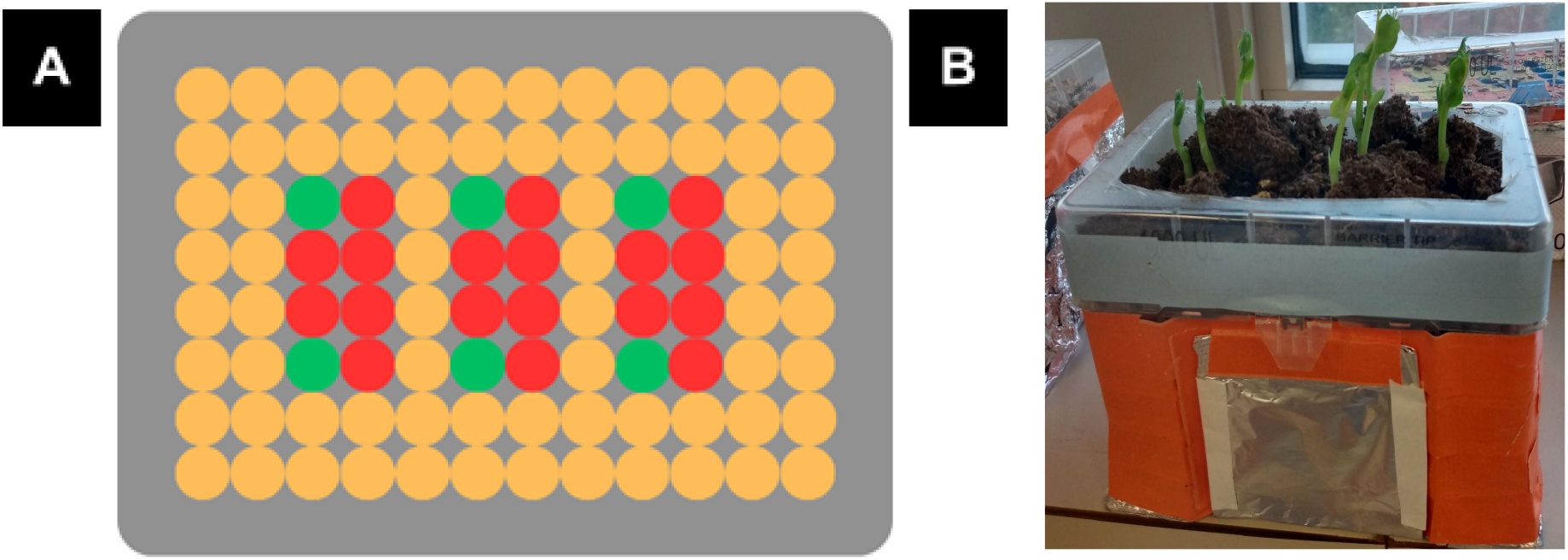
Visual Representation of the Hydroponic Apparatus. Figure 1 [A] is a schematic showing the layout of the seed tray used in the isolation device. Orange represents holes covered with laboratory tape, green represents holes pea seedlings were placed on, and red represents holes that had the plastic connecting them removed. Figure 1 [B] shows a fully completed device.

To begin seed growth: six AC Carver pea seeds were allowed to incubate in a 250mL flask filled with 30mL of sterile water overnight (approximately 24 hours). Then, 1.5mL of stock nutrient solution (1.00 M Potassium Nitrate, 0.255 M Calcium Phosphate, 0.564 M Magnesium Sulfate, 0.130 M Ammonium Sulfate, and 8.16 mM Ferric Citrate) was added to 500mL of sterile water and placed into the base of the hydroponic system by removing the seed tray.^4^ Aiding in the initiation of the experiment, the stock nutrient solution was found to be stable at 4°C for at least three months. 2g of soil infected with *A. euteiches* and a stir bar were added, and the seed tray was placed into the system. Six pre-soaked AC Carver Pea seeds were then added to their appropriate positions on the seed tray, as shown in Figure 1. To increase efficiency, roots of germinating seeds were placed pointing into the nutrient solution when applicable. Peat Moss broken into pieces measuring approximately 2 cm^3^ was then added on top of empty gaps in the seed tray. Small grain vermiculite was then used to fill the upper chamber of the system, but this was found to be purely cosmetic and had no effect on seed growth. Sterile water (approximately 100mL) was then slowly added to the top of the system until the peat moss and vermiculite were wet. The system was then placed by a window on a stirring plate set at low. Because the experiment occurred during the summer, light conditions were satisfactory. After two days, the stirring plate was removed and the plants were allowed to grow without further intervention. Isolates were identified by plating infected roots on MVB plates and subculturing potential isolates onto EMD potato dextrose agar or MilliporeSigma corn meal agar.^3^ The species of isolate was confirmed by amplifying the CPN60 UT with site-specific primers (unpublished) and sequencing the amplicon using Sanger DNA sequencing.

### Isolate Pea Trial

A square of agarose from a culture containing *A. euteiches* deemed non-viable by the typical culture propagation method was placed into a 50 mL falcon tube containing 25 mL EMD potato dextrose broth. This was then allowed to incubate in a shaking incubator for approximately a week at 20°C and then allowed to incubate at room temperature for an additional three weeks. 12.5 mL of this was then used to inoculate vermiculite-filled pots containing an AC Carver pea seed. When the plants grew they were transplanted into a bigger pot, transferring the vermiculite from the previous pot with the plant. After harvesting, DNA was extracted from the roots using a Quiagen Plant Pro Kit. The roots of the plant infected with an isolate from the previous step, hereafter called Carefoot #3 were also plated for isolation on MVB as described by Pfender et al.^3^

### Molecular Diagnostics

The *Aphanomyces* CPN60 LAMP assay was performed using Optigene’s ISO-001 LAMP Isothermal Master Mix using an unpublished assay dubbed AE CPN60 LAMP. qPCR was performed using the assay developed by Chatterson et al. or an unpublished variant of it.^5^

## RESULTS

### Effect of Hydroponic Design

Throughout the experiment, the hydroponic apparatus was manipulated with the hopes of increasing the efficiency at which *Aphanomyces euteiches* could be isolated. Through this process, it was found that *A. euteiches* can not infect plants unless a stir bar is initially present. This may be due to its inability to reach the nascent pea roots. It was also found that if a stir bar was left spinning for too long, damage to the roots of the pea seedlings could occur, reducing the yield of culturable material. Seed germination time was found to be a bottleneck for culturing, as it varies between individuals. Initially, seeds were placed into the hydroponic apparatus without pre-soaking. This resulted in an unsatisfactorily slow germination rate (up to four to five days) and necessitated that the seed be held in place, as it was initially too small to fit into the seed tray. Pre-soaking seeds drastically increased the germination rate, in many cases to as little as one day. It also allowed peas to germinate within a narrower timeframe, allowing a more accurate measurement of when they should be harvested for culturing. Twenty-four hours was found to be the optimal time for soaking the seeds. Longer soaks were found to promote the growth of microorganisms that could stunt later seed development.

### Findings of Culturing

Because each seed was unique, there was no ideal day to harvest the pea roots for plating. However, the optimal time for attempting to culture was found to be around six days after pdescribed in the material and methods section, up to seven unique *A. euteiches* individuals can lacement into the apparatus, when the plants had a solid taproot with 4-6 developing lateral roots. At this time, the roots of the seedlings would appear mostly white, with a few patches of minor browning. If the seedling was allowed to continue to grow, the roots would become a deep honey brown, as shown in Figure 2 [A]. Highly infected roots such as this were found to be not ideal for culturing *A. euteiches* due to the presence of other pathogens that could outcompete *A. euteiches* on the MVB plate, including *Rhizopus*. Using the procedure be harvested per hydroponic apparatus. Two days after plating was found to be the optimal time for subculturing. To test the effectiveness of the apparatus, five suspected *Aphanomyces* isolates were selected for CPN60 sequencing. All five of these had a 100% match to the GenBank sequence CU365658.1 from base pair 411 to 752. One strain, Carefoot 11, was also assayed using the *Aphanomyces* qPCR described by Shayma et al.^5^

**Fig. 2.**
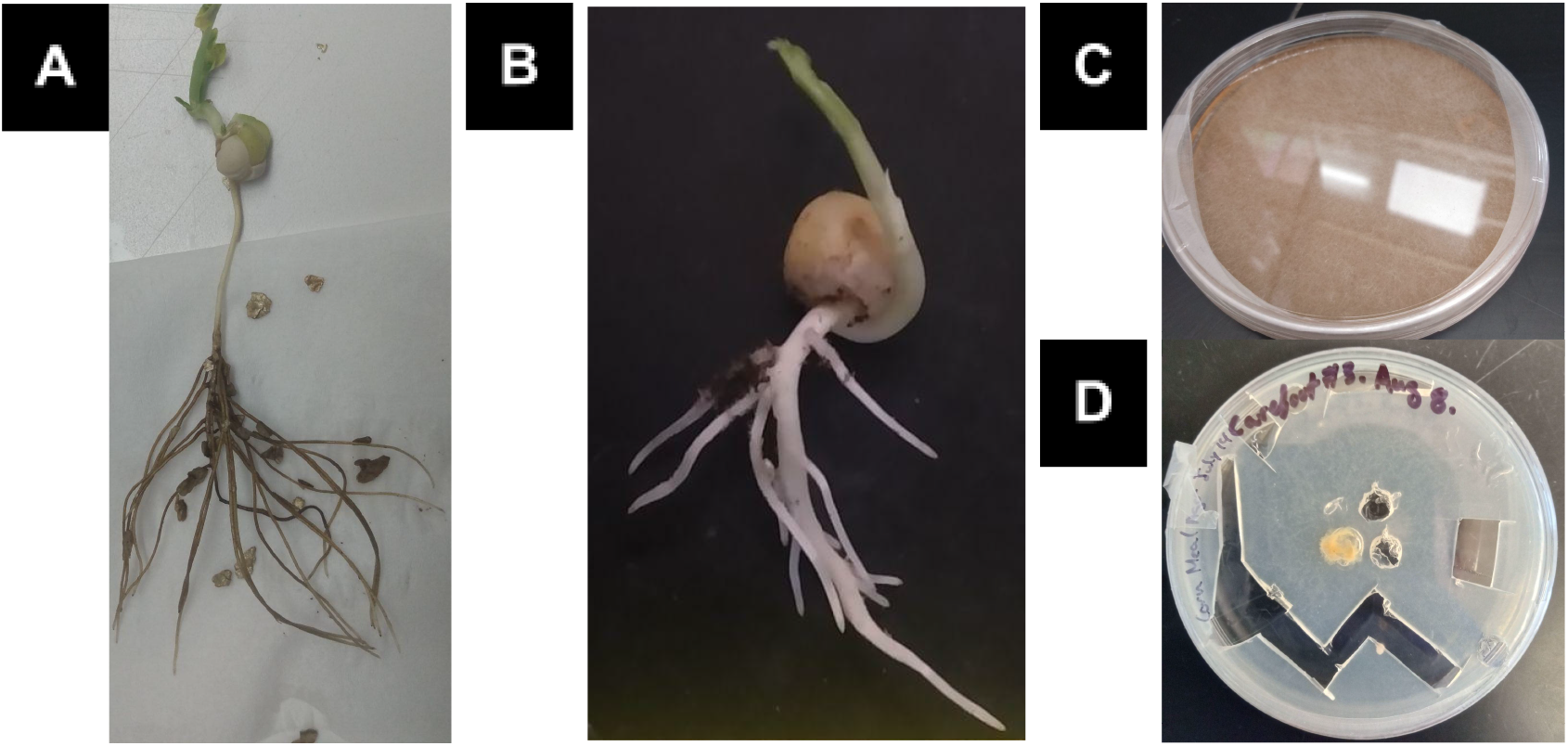
Products of Hydroponic Culturing. Figure 2 [A] An infected pea root that did not yield viable isolates. Figure 2 [B] An infected pea root harvested on day six which yielded a potential Aphanomyces euteiches isolate. Figure 2 [C] A confirmed A. *euteiches* isolate (Carefoot #11) from the hydroponic apparatus with typical arial mycelia. Figure 2 [D] The Carefoot #3 *A. euteiches* isolate lacking aerial mycelia.

Despite this similarity, two of the isolates, arbitrarily called Carefoot #1 and Carefoot #3, shown in Figure 2 [D], displayed unique slow-growing phenotypes. Upon further sequencing, these contained the correct CPN60 sequence but had a similar, yet unique, ITS sequence. Unfortunately, these isolates stopped growing using the typical propagation method. In an attempt to revive these isolates, they were inoculated into plants without negative or positive controls. However, surprisingly, as shown in Figure 4 these isolates not only caused an infection, as evidenced by an unpublished *A. euteiches* Taq-man qPCR (Cq 26.40 - 30.39), but also allowed the plants to reach maturity as shown in Figure 3. This is especially surprising, as the soil they originated from was lethal to AC carver peas within two weeks of planting (unpublished data). Upon culturing from the plants, these isolates returned positive for *A. euteiches*. However, as shown in Table 1, they had a different annealing temperature compared to the classical isolate (Carefoot #11). Future research will focus on elucidating this isolate.

**Table 1:** CPN60 LAMP Assay Results for Carefoot #3 Plant Isolates.

| Sample Name | Amplification Time (minutes:seconds) | Annealing Temperature (°C) |
| --- | --- | --- |
| Isolate 3.P1 | 21:30 | 86.91 |
| Isolate 3.P2 | 28:15 | 86.67 |
| Isolate 3.P3 | 39:00 | 86.57 |
| Carefoot 11 | 16:30 | 85.77 |
| Water | Na | 85.92 |

**Fig. 3.**
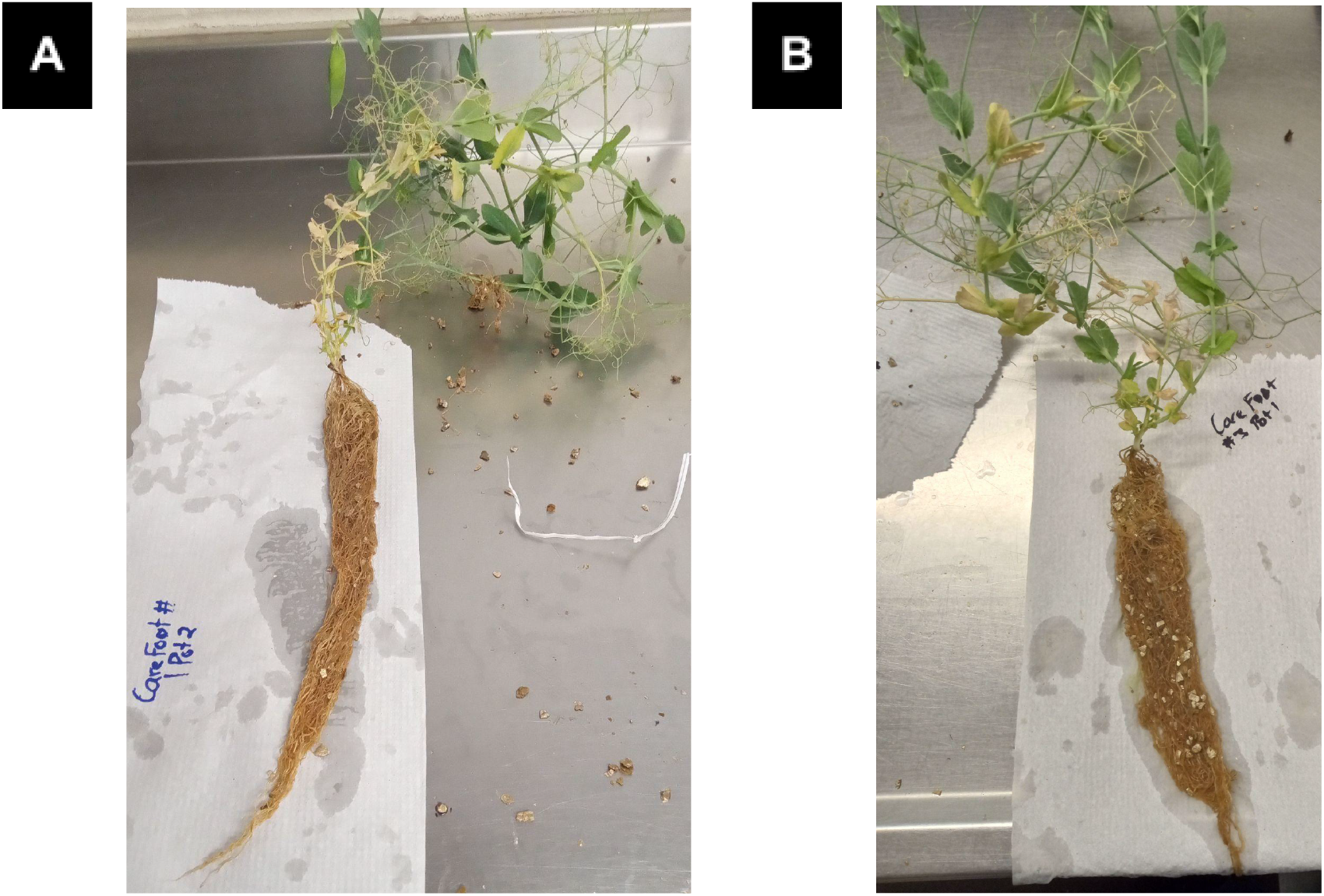
Experimental Aphanomyces Plants. Figure 3 [A] The plant infected with Carefoot 1 had a Cq of 26.66. Figure 3 [B] The plant infected with Carefoot 3 had a Cq of 26.40. These were determined using the unpublished qPCR assay. Further washing did not remove the coloration of the roots.

**Fig. 4.**
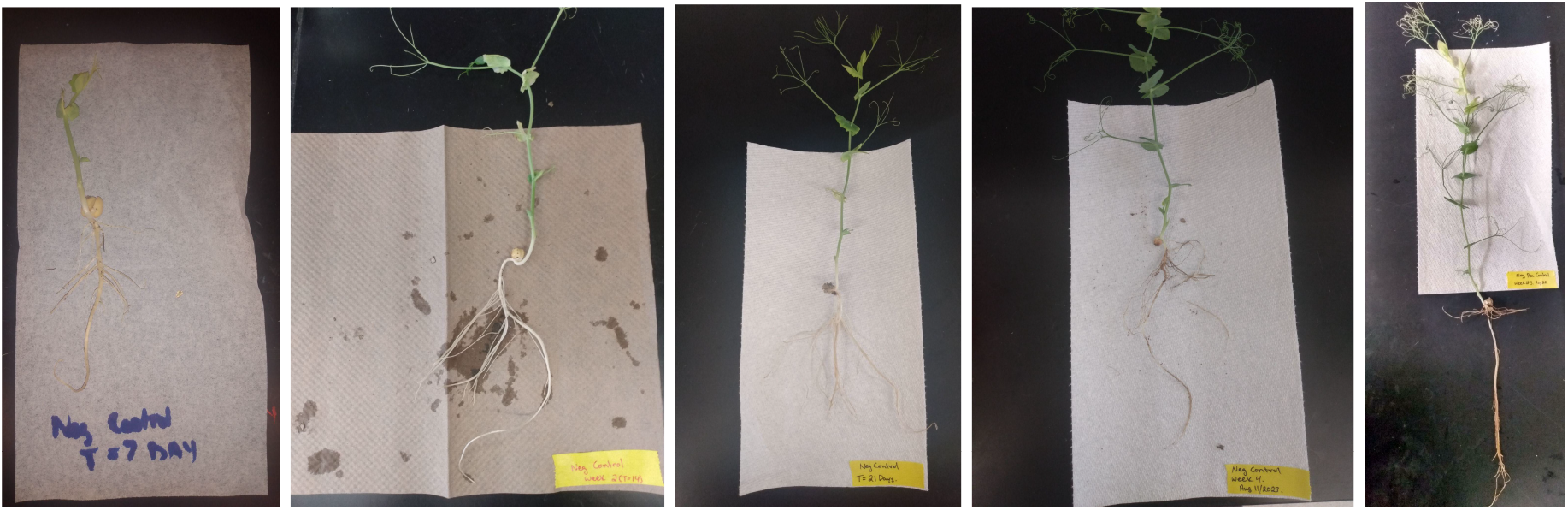
Timeline of Negative Control. Timeline of the negative control experiment over five weeks. Note the whiteness of the roots.

**Fig. 5.**
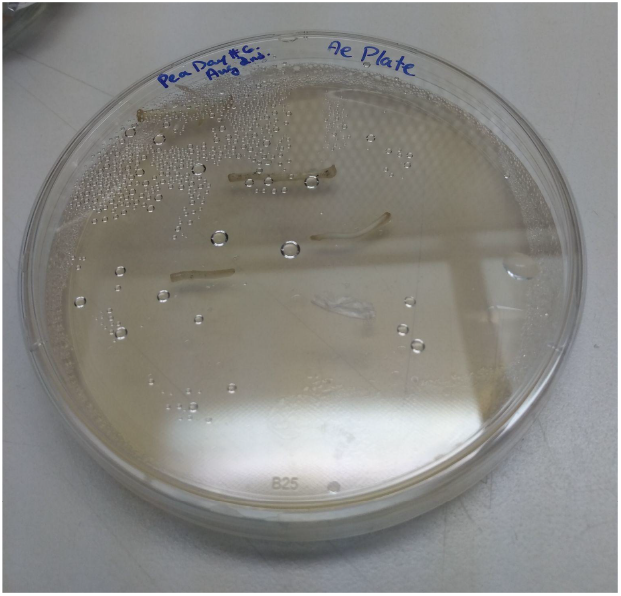
Plating of Infected Roots on MVB Media. Roots were plated on MVB media containing antibiotics.

### Limitations of the Growth Apparatus

When infected soil was not added, the pea plants could grow for up to five weeks so long as nutrient solution was continually added when depleted. Uninfected pea plants had no discoloration on their roots until four weeks after planting. At this time, leaf discoloration also began to appear. At week five, further discoloration of the roots was visible, and more leaves began to turn yellow. It is unknown why this occurred. Figure 4 shows the stages of pea development in the absence of infected soil.

## DISCUSSION

The results indicate that when soil known to contain *Aphanomyces euteiches* spores is added to a hydroponic system containing a susceptible host, infection will occur. Infection is severe enough that infected host root tissue can then be used to isolate *A. euteiches* on an MVB plate.^3^ Plants grown in the hydroponic system were found to be much more efficient in creating opportunities to isolate *A. euteiches* than those grown in soil, in part due to the ability to view the growth progress of each plant and harvest it at the ideal time, but also likely due to the lack of contaminated soil clinging to the hydroponically grown plants. Another issue with culturing *A. euteiches* this hydroponic apparatus may ameliorate is its ability to survive storage. Currently, *A. euteiches* spores do not store well in artificial media. However, they are known to last up to ten years in field soil.^1^ The ability to easily isolate *A. euteiches* from soil may present a new opportunity for storage. Future research may consider how sterile soil can be inoculated with oospores of a particular culture, stored, revived using the hydroponic system, and then isolated from the resulting infected pea roots.

But what was most surprising about this experiment is that throughout this project, two strange isolates were found that had a unique translucent branching structure and that did not produce aerial mycelia. Research is ongoing to determine what the identity of these microorganisms is. This may be a phenotype associated with a mixed culture. However, should these isolates truly prove to be *A. euteiches*, it may indicate that this methodology can be used to find the existence of rare *A. euteiches* strains that are otherwise difficult to phenotype. Given that plants infected with these managed to produce a yield, these strains may also present a future biocontrol opportunity. It is important to note, however, that the plant trial was not intended as an infection trial, only a culture revival. As such, this evidence may be an experimental artifact as no negative controls were used. However, given how little is known about *A. euteiches*, it should motivate future work focusing on elucidating the phenotypic diversity of *A. euteiches*.

I confirm that I do not have a conflict of interest.

