## Supplementary figures and images for "Hydroponics: A Novel Approach to Culturing *Aphanomyces euteiches*"

### Supplemental Figure 2

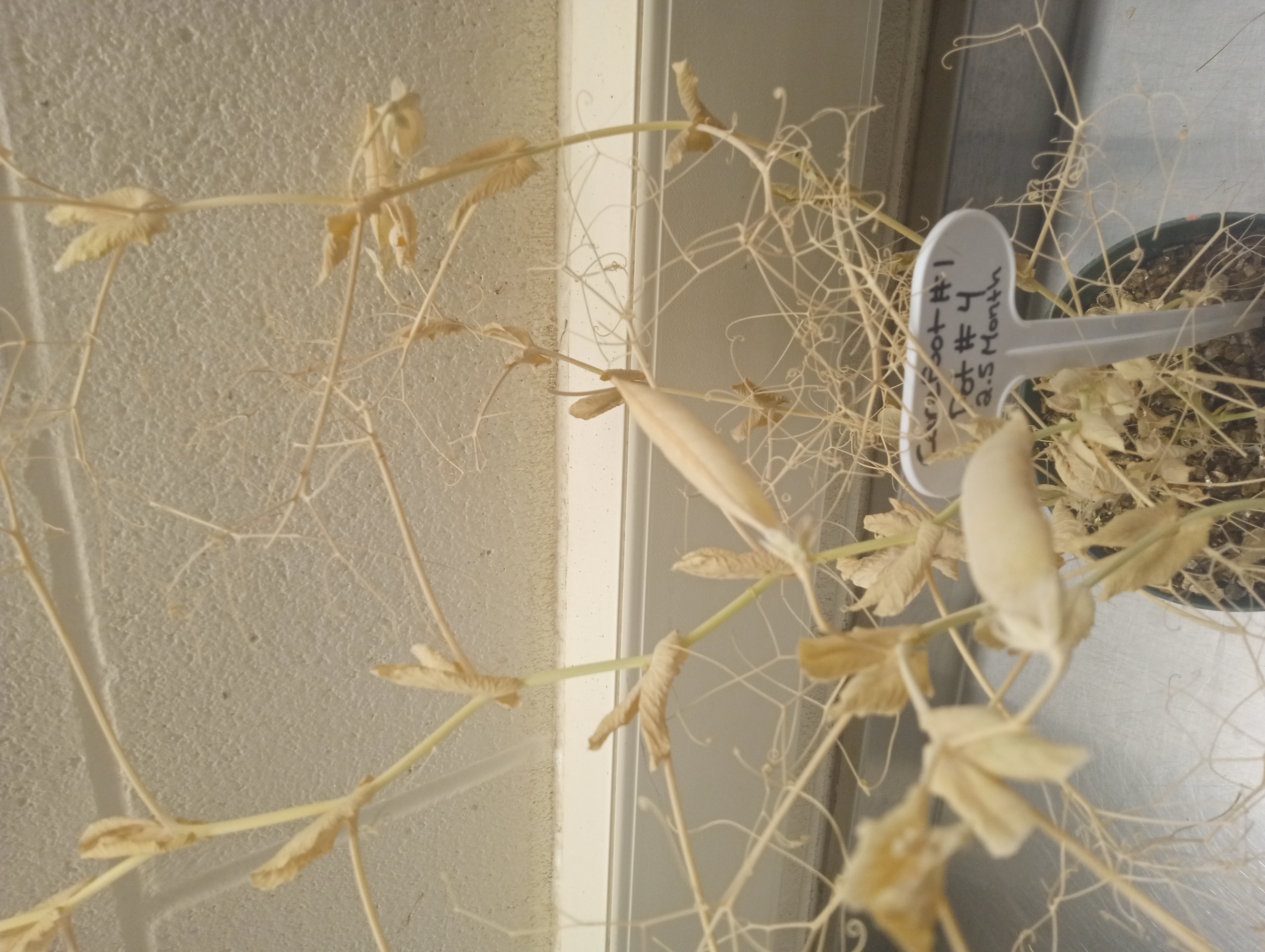
